# Fluorescence *In Situ* Hybridization Provides Evidence for the Presence of the Endosymbiotic Bacterial Genus *Rickettsia* in Tardigrades

**DOI:** 10.1101/2022.12.01.518763

**Authors:** Bienvenido W. Tibbs-Cortes, Dylan L. Schultz, Laura E. Tibbs-Cortes, Stephan Schmitz-Esser

**Affiliations:** Interdepartmental Microbiology Program, Iowa State University, USA; Genetics and Genomics Program, Iowa State University, USA

**Keywords:** Endosymbiont, Fluorescence *In Situ* Hybridization, Microbiota, *Rickettsia*, Tardigrades

## Abstract

Many ecdysozoans harbor endosymbiotic bacteria within their microbiota, and these endosymbionts can have a range of positive and negative effects on their hosts. Recent 16S rRNA gene amplicon sequencing studies have provided evidence for endosymbionts within the tardigrade microbiota. In a previous study amplicon study, we determined that sequences corresponding to the endosymbiotic genus *Rickettsia* were significantly more associated with tardigrades than with the substrate from which they were isolated. Here, we performed fluorescence *in situ* hybridization (FISH) using a *Rickettsia*-specific probe, RickB1, to determine if *Rickettsia* could be found in tardigrades. RickB1 and a probe targeting most bacteria, EUB338, colocalized within the tissues of tardigrades, indicating the presence of *Rickettsia*. We also performed FISH using RickB1 and a nonsense probe which allowed us to distinguish between false positives and true positives. This method revealed RickB1 signals in tardigrades that were not due to erroneous probe binding, providing further evidence that *Rickettsia* is present in tardigrades. Future research will be necessary to determine the effects, if any, of this endosymbiont on its tardigrade host.

## Introduction

Tardigrades are a globally ubiquitous phylum of microscopic ecdysozoans well known for their ability to survive extreme conditions, but knowledge of the tardigrade microbiota is relatively limited. In general, the microbiota, often used synonymously with microbiome or microbial community, is made up of the microorganisms present in a given environment (Eisenhofer et al., 2019; Mushegian et al., 2019; Ursell et al., 2012). In the case of other ecdysozoan animals including *Caenorhabditis elegans, Daphnia magna*, and various insects of multiple orders, members of the microbiota may be acquired horizontally from the environment, food, and social interactions as well as vertically from mother to offspring (Engel & Moran, 2013; Mushegian et al., 2019; Zhang et al., 2017). Some of these microorganisms will be transient members of the microbial community, while others will colonize the animal and persist over long time periods (Engel & Moran, 2013; Mushegian et al., 2019; Zhang et al., 2017). This differential persistence of microorganisms in ecdysozoan microbiota can be due to factors including host genetics, behavior, and immune function as well as inter-species competition (Mushegian et al., 2019; Portal-Celhay & Blaser, 2012). Identifying the sources of inoculum for the tardigrade microbiota as well as characterizing transient and persistent microorganisms associated with tardigrades is an ongoing subject of research.

The first studies of tardigrade-associated bacteria, conducted in 1999 and 2000, found plant pathogens associated with *Macrobiotus hufelandi* C.A.S. Schultze, 1834 and a non-random association between tardigrades and bacteria (Benoit et al., 2000; Krantz et al., 1999). More recent studies have used 16S rRNA gene amplicon sequencing to survey the tardigrade microbiome in at least 13 tardigrade species collected from locations in Antarctica, Curaçao, Ecuador, Madagascar, and the United States, as well as in Italy, Poland, Sweden, and the United Kingdom. These surveys found that tardigrade microbiota are altered by laboratory culture and are distinct from and usually less diverse than the microbiota of their substrate (Boscaro et al., 2022; Kaczmarek et al., 2020; Tibbs-Cortes et al., 2022; Vecchi et al., 2018; Zawierucha et al., 2022). *Proteobacteria* has been consistently reported as the most abundant bacterial phylum associated with tardigrades, while *Firmicutes, Bacteroidetes*, and *Actinobacteria* have each been reported as highly abundant in at least three previous surveys of the tardigrade microbiota (Kaczmarek et al., 2020; Mioduchowska et al., 2021; Tibbs-Cortes et al., 2022; Vecchi et al., 2018).

The presence of endosymbiotic bacteria in the tardigrade microbiota is also an ongoing area of research. Endosymbiotic bacteria survive within host cells, and relationships between endosymbiont and host range from pathogenic to mutualistic (McCutcheon et al., 2019; Taylor et al., 2012). Because of their intracellular nature, endosymbionts of invertebrates are often transmitted vertically from mother to offspring although horizontal transmission is known to occur (McCutcheon et al., 2019; Moran et al., 2008). Notably, the recent 16S rRNA gene amplicon sequencing studies of tardigrade microbiota have provided evidence for the existence of endosymbiotic bacteria in tardigrades (Guidetti et al., 2019b; Kaczmarek et al., 2020; Mioduchowska et al., 2021; Vecchi et al., 2018). Vecchi et al. (2018) identified operational taxonomic units (OTUs) corresponding to members of the obligate endosymbiotic order *Rickettsiales* in the microbiota of six tardigrade species. Subsequent diagnostic PCRs of tardigrades extracted from the same substrates provided additional evidence for the existence of *Rickettsiales* in tardigrades and further identified the *Rickettsiales* OTUs as members of *Anaplasmataceae* and *Ca. Tenuibacteraceae* (Guidetti et al., 2019b). Importantly, the same study by Guidetti et al. (2019b) also performed fluorescence *in situ* hybridization (FISH) and observed bacteria within the body cavity of *Echiniscus trisetosus* Cuénot, 1932. Similarly, more recent 16S rRNA gene amplicon sequencing studies by Kaczmarek et al. (2020) and Mioduchowska et al. (2021) identified putative endosymbionts in tardigrades. However, these studies did not sequence negative control samples to account for potential contaminants or conduct additional follow-up assays to confirm their results (Kaczmarek et al., 2020; Mioduchowska et al., 2021).

The recent analysis by our group also yielded sequencing-based evidence suggesting the presence of endosymbionts in tardigrades. Specifically, an OTU (OTU 180) classified as a member of the genus *Rickettsia* was significantly more abundant in the tardigrade community microbiota than in the substrate (Tibbs-Cortes et al., 2022). *Rickettsia* is a diverse genus consisting of obligate intracellular symbionts with a wide variety of hosts including mammals, leeches, cnidarians, and insects (Davison et al., 2022; McGinn & Lamason, 2021). Many *Rickettsia* species are capable of manipulating the reproduction of their hosts through male-killing and the induction of parthenogenesis (Hagimori et al., 2006; Massey & Newton, 2022; von der Schulenburg et al., 2001). Indeed, the presence of *Rickettsia* in a host is often associated with a decrease in fecundity, but in other instances, infection with *Rickettsia* has been ocumented to impart beneficial effects on the host including resistance to thermal stress and fungal pathogens (Brumin et al., 2011; Łukasik et al., 2013; McGinn & Lamason, 2021).

16S rRNA gene amplicon sequencing is a useful tool to survey the community structure and composition of tardigrade microbiota; however, because it reports only the presence of DNA, additional evidence from other methods will be necessary to confirm the presence of cells of taxa of interest as well as their localization. Here, we utilized FISH to confirm the presence of *Rickettsia* in tardigrades from than same substrate samples used in our previous study. We identified signals likely belonging to *Rickettsia* within these tardigrades using confocal microscopy, providing additional evidence that some tardigrades harbor endosymbiotic bacteria.

## Methods

### Tardigrade Isolation

We previously analyzed the microbiota of tardigrade communities collected from apple orchards in central Iowa, USA. The OTU classified as *Rickettsia* was most abundant in the tardigrades from lichen samples collected in 2020 at Location 1 (Tibbs-Cortes et al., 2022).

Therefore, we extracted additional tardigrades from these samples for this study. Lichen samples were soaked overnight in water purified using a Milli-Q system (EMD Millipore). Subsamples of water were collected from the rehydrated lichen, and tardigrades were isolated from these with Irwin loops using a dissecting microscope. Tardigrades were placed in drops of PCR-grade water, and the Irwin loop was sterilized using a flame after each tardigrade. Tardigrades were washed by transferring to a fresh drop of PCR-grade water, and this process was repeated for a total of three washes. All tardigrade isolation steps were carried out in a sterile field created by a Bunsen burner flame.

### Probes

The FISH probes used in this study are listed in Table 1. EUB338 is a general probe that targets the 16S subunit of ribosomal RNA of the majority of bacteria species (Amann et al., 1990). NONEUB is a nonsense probe with a sequence complementary to that of EUB338.

**Table 1.**
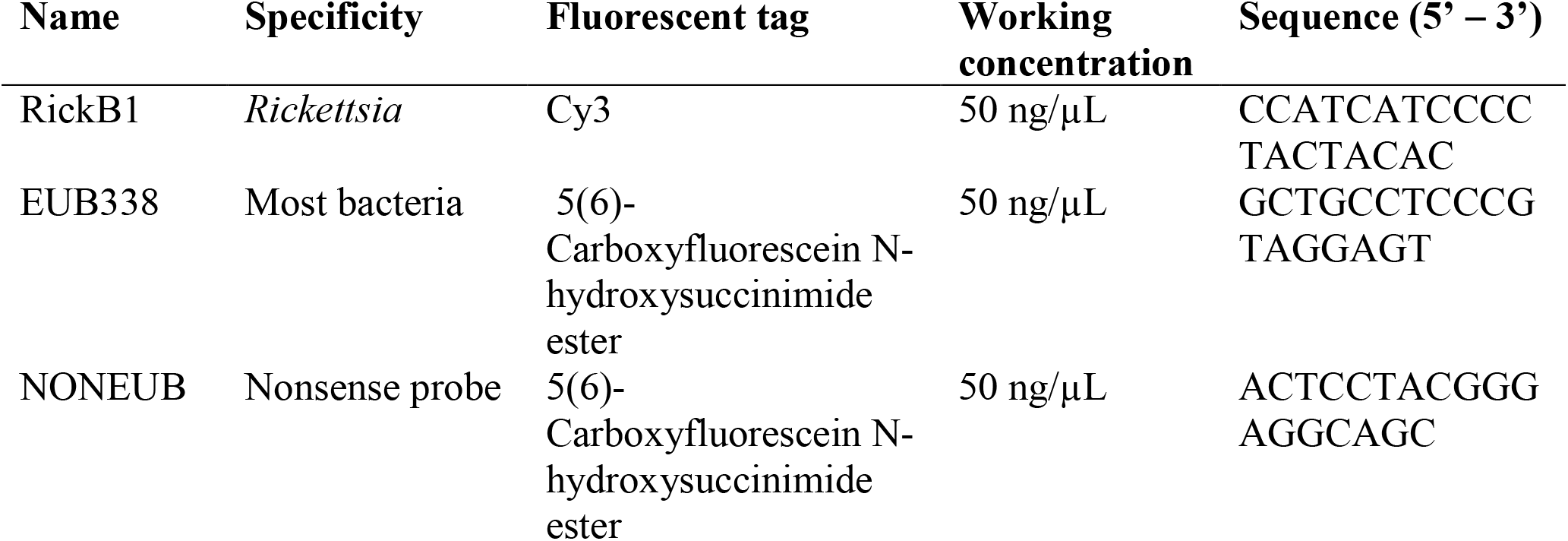
Probe information

Because it does not bind to the ribosomal RNA of any bacteria, it is used to identify false positives such as those caused by erroneous probe binding (Manz et al., 1992). Finally, the probe RickB1 targets members of the genus *Rickettsia* (Perotti et al., 2006). Samples were hybridized with EUB338 and RickB1 to identify any bacteria within tardigrades as well as to identify bacteria belonging to the genus *Rickettsia*. Staining with NONEUB and RickB1 allowed a more stringent identification of *Rickettsia*, as colocalization of the two probes would indicate a false positive.

### Fluorescence *in situ* hybridization

FISH was performed using a protocol modified from Daims et al. (2005). Washed tardigrades were dried onto one field of a 76 × 26 × 1 mm 10-well (6mm) polytetrafluoroethylene-printed slide (Electron Microscopy Sciences). Fixation of specimens was performed by adding 30 μL of ice-cold 3:1 4% paraformaldehyde:1x phosphate buffered saline (PBS) to the field. The slide was placed in a sterile petri dish sealed with parafilm, and the specimens were fixed at 4°C for 2 hours. The paraformaldehyde solution was then removed with a pipette, and tardigrades were washed three times with ice-cold 1x PBS. Specimens were allowed to dry on the slide. An ethanol dehydration series was conducted by applying 30 μL of 50% ethanol to the specimens. After 3 minutes, the ethanol solution was removed using a pipette. This was repeated with an 80% ethanol solution and then with 100% ethanol. Fixed and dehydrated tardigrades were dried on the slide.

After drying, 10 μL of 35% formamide hybridization buffer (Table 2) was pipetted onto the slide fields containing fixed tardigrades. The web tool mathFISH was used to determine the appropriate formamide concentration (Yilmaz et al., 2011). 1 μL of probe RickB1 and 1 μL of either probe EUB-338 or probe NONEUB were then pipetted directly into the hybridization buffer. No-probe staining of specimens was also performed as a control. A humid hybridization chamber was prepared by placing a small piece of paper towel soaked with 35% formamide hybridization buffer into a 50mL screw-cap tube. The slide was then placed inside the hybridization chamber with the cap screwed shut, and the chamber was incubated horizontally at 46°C for 1.5 hours. Following hybridization, the hybridization buffer was removed from the samples via pipetting, and tardigrades were rinsed once with 35% formamide wash buffer (Table 2) pre-warmed to 48°C. Next, another 30 μL of the pre-warmed wash buffer was pipetted onto the samples. A humid wash chamber was created as above using 35% formamide wash buffer to soak the paper towel. The slide was then incubated inside for 10 minutes at 48°C. Finally, the wash buffer was removed, and the samples were washed twice with ice-cold MilliQ water and dried completely. Citifluor™ AF1 antifadent solution (Electron Microscopy Sciences) was added to the fields, and a cover slip was placed over the specimens. The cover slip was sealed onto the slide using clear nail polish, and the slide was stored in the dark at 4°C until visualization.

**Table 2.** Composition of hybridization buffer and wash buffer solutions.

|  | 35% formamide hybridization<br>buffer (1 mL total volume) | 35% formamide wash buffer (50<br>mL total volume) |
| --- | --- | --- |
| MilliQ H<sub>2</sub>O | 449 μL | 47.8 mL |
| 5 M NaCl | 180 μL (900 mM) | 0.7 mL (70 mM) |
| 1 M Tris-HCl | 20 μL (20 mM) | 1 mL (20 mM) |
| Formamide | 350 μL (35%) | 0 mL |
| 10% SDS | 1 μL (0.01%) | 0 mL |
| 0.5 M EDTA | 0 μL | 0.5 mL (5 mM) |

### Microscopy

Preliminary FISH experiments using probes RickB1 and EUB338 were visualized with epifluorescence and phase contrast microscopy using a Leica DM4B upright microscope with a Leica N PLAN 40x/0.65 dry objective. For fluorescence microscopy, an LED and two filter cubes were used for excitation of fluorophores and detection of emission, respectively. The filter cube used to visualize localization of RickB1 allowed transmission of light emitted at 605 nm, and the filter cube used to visualize EUB338 allowed transmission of light at 527 nm. Fluorescence and phase contrast images were taken with the mounted Leica DFC3000 G microscope camera using the Leica Application Suite × (LAS X) software.

Confocal microscopy using a Leica Stellaris STED microscope with a Leica HC PL APO CS2 40x/1.30 oil immersion objective was later used to visualize additional specimens stained with probes RickB1 and NONEUB. Excitation of RickB1 was conducted with a 555 nm laser with 2.05% intensity, and NONEUB was excited with a 494 nm laser at 6.19% intensity. Two HyD X detectors set to detect 561-675 nm and 500-550 nm were used to detect the emissions of RickB1 and NONEUB, respectively. Line averaging was performed with a total of four scans, and differential interference contrast (DIC) images were acquired simultaneously. Images were captured using LAS X.

## Results

In initial experiments, tardigrades were stained with both the *Rickettsia*-specific probe RickB1 and the general bacteria probe EUB338 to identify bacteria and *Rickettsia* in tardigrade tissues. Epifluorescence microscopy revealed that RickB1 and EUB338 colocalized to circular points within the body cavity in a specimen of *Paramacrobiotus tonollii* (Ramazzotti, 1956) (Figure 1). These points were approximately 1-2 μM in diameter and were primarily located within the dorsal region between legs III and IV. Additional points in this region of the body cavity were observed with EUB338 signals but not RickB1 signals (Figure 1, white arrows). In this specimen, no instances of RickB1 colocalization were observed without simultaneous colocalization of EUB338. Additionally, no features similar to the points displayed in Figure 1 were observed in the no-probe controls. Although other specimens were stained with the two probes, the strong levels of autofluorescence made it difficult to examine these other specimens for RickB1 signals.

**Figure 1.**
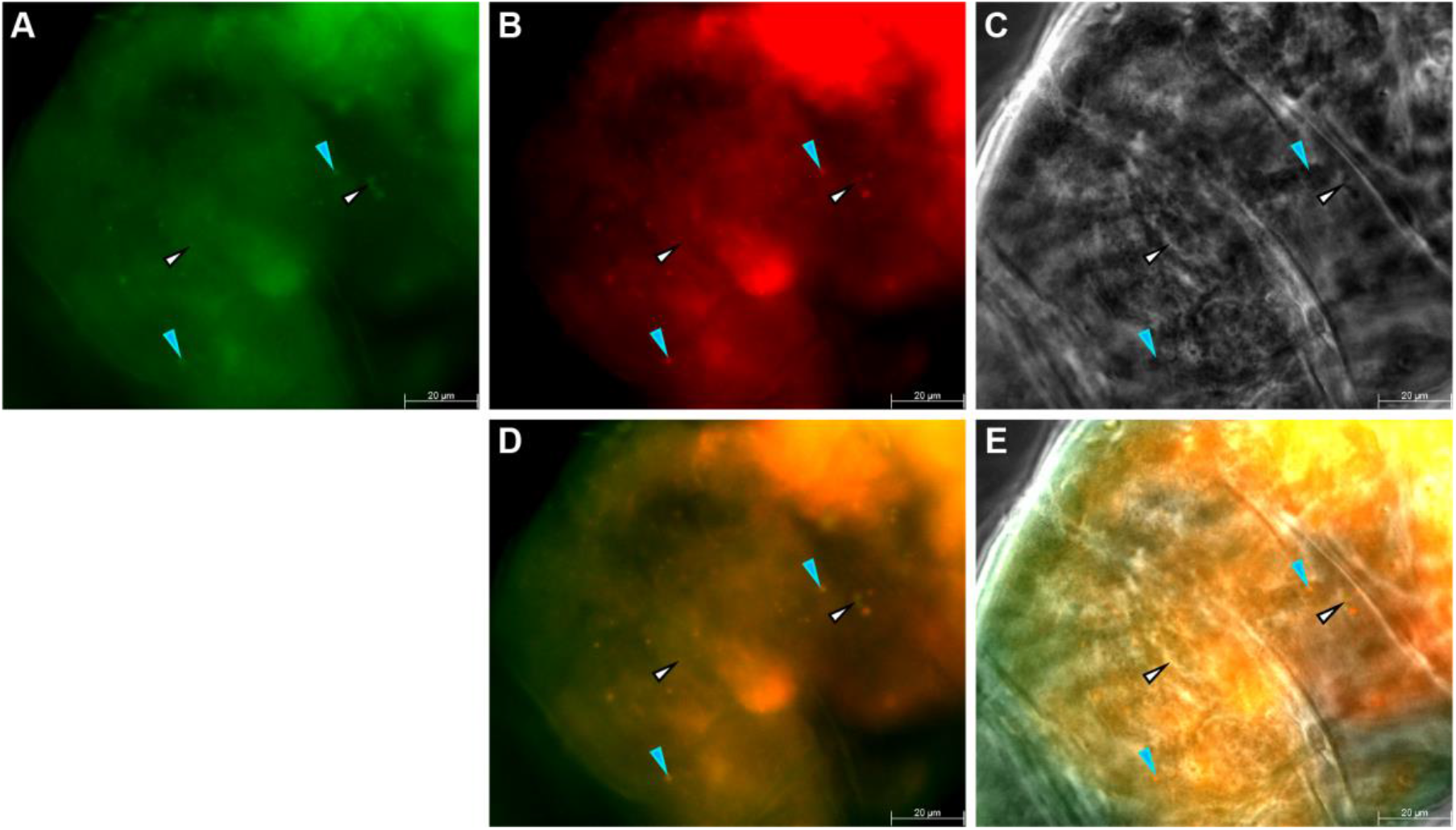
Epifluorescence microscopy reveals that RickB1 and EUB338 colocalize at points within the body cavity of a specimen of *Paramactobiotus tonollii*. The viewing field is focused on the dorsal side of the animal between legs III and IV. Blue arrows point to features with colocalization of EUB338 and RickB1, likely *Rickettsia*. White arrows point to features with only an EUB338 signal, possibly another bacterium not belonging to *Rickettsia*. Scale bars = 20 μm. **(A)** Green channel for detection of EUB338 **(B)** Red channel for detection of RickB1 **(C)** Phase contrast **(D)** Overlay of green and red channels. Overlap of green and red indicates colocalization of EUB338 and RickB1 (orange) **(E)** Overlay of fluorescence channels and phase contrast.

Subsequent experiments utilized RickB1 and the nonsense probe NONEUB to identify *Rickettsia* within tardigrades and simultaneously detect false positives. Due to the high levels of autofluorescence from tardigrade tissues, we utilized confocal microscopy to analyze samples labeled with RickB1 and NONEUB. Confocal microscopy revealed that in some specimens, such as the simplex *Macrobiotidae* seen in Figure 2, RickB1 and NONEUB colocalized, indicating a false positive for *Rickettsia*. In some of these cases the signals of the two probes corresponded to cuticular structures that were visible with DIC microscopy (Figure 2D, white arrow). The no-probe controls did not exhibit signals comparable to those shown in Figure 2.

**Figure 2.**
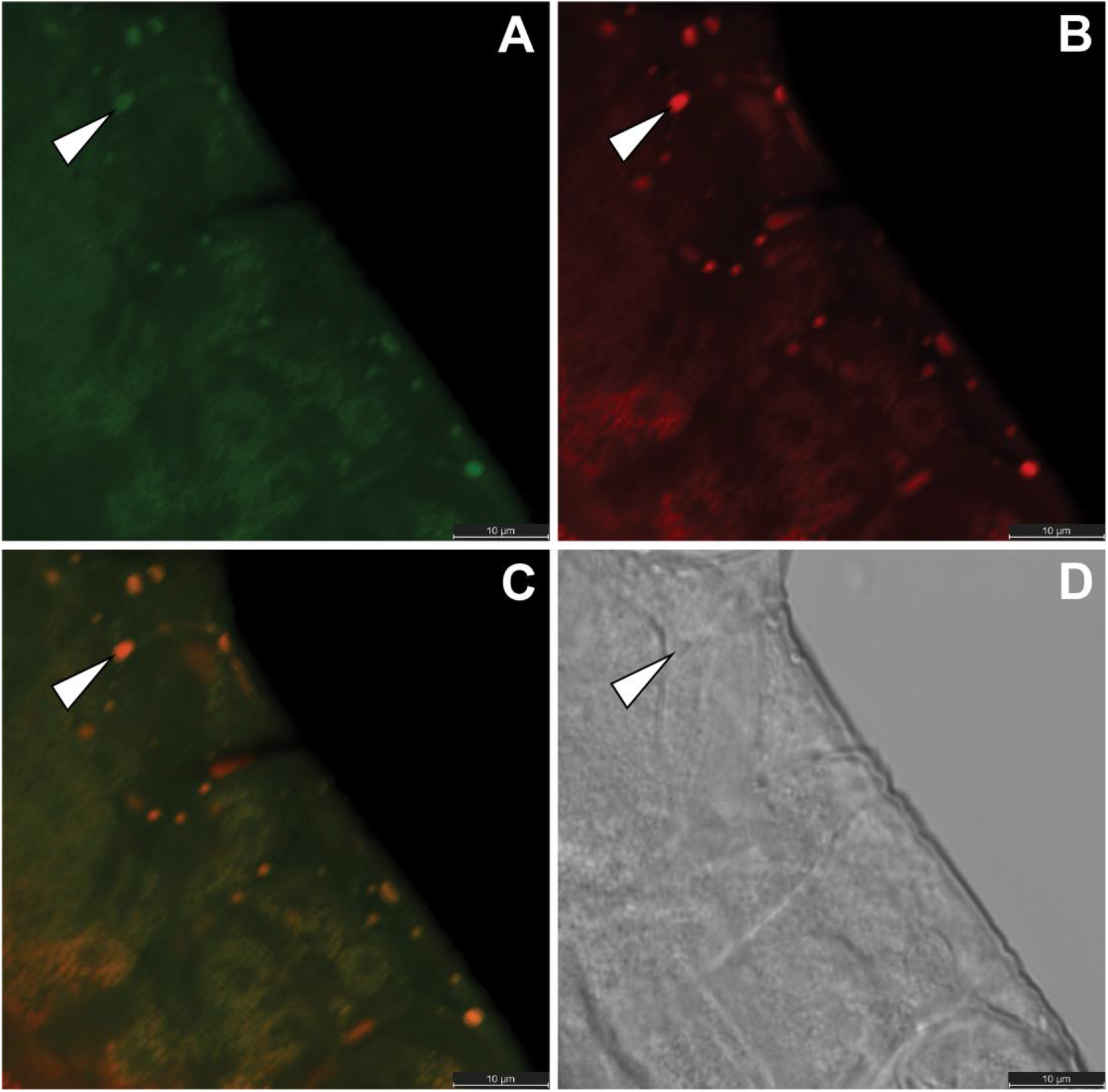
Labeling with NONEUB allows detection of false positives due to erroneous probe localization. Confocal microscopy was used to visualize a simplex *Macrobiotidae* sp. labeled with RickB1 and NONEUB. The NONEUB probe sequence will not bind to the ribosomal RNA of any bacteria. Therefore, colocalization of NONEUB with RickB1 indicates a false positive due to erroneous probe localization or autofluorescence. The field shows the lateral side of the specimen between legs I and II. White arrow points to one region of NONEUB and RickB1 colocalization that is also visible as a structure in DIC. Scale bars = 10 μM, pixel size = 0.037 by 0.037 μM, zoom = 3.83. **(A)** Multiple areas of NONEUB signal (green) are visible throughout the specimen **B)** Multiple RickB1 signals corresponding to the NONEUB signal are present **(C)** Overlay confirms the colocalization of RickB1 and NONEUB (orange), demonstrating that NONEUB can be used to identify erroneous signals that would lead to false positives **(D)** Certain features of the specimen visible through DIC correspond to regions of NONEUB and RickB1 colocalization.

However, one specimen of *Paramacrobiotus* was observed that possessed multiple RickB1 signals without colocalization of NONEUB (Figure 3). These areas of colocalization were slightly less than 1 μm in diameter, somewhat smaller in size than those observed in the *P. tonollii* specimen in Figure 1. These signals were also smaller in diameter than the false positive signals observed in the simplex *Macrobiotidae* specimen (Figure 2). These points were located within the head of the animal and did not correspond to any structures visible with DIC. Due to positioning of the specimen on the slide, it was not possible to view the rest of the body to determine if additional signals were seen in the rest of the specimen. The no-probe controls did not exhibit signals similar to those in the *Paramacrobiotus* sp. (Figure 3).

**Figure 3.**
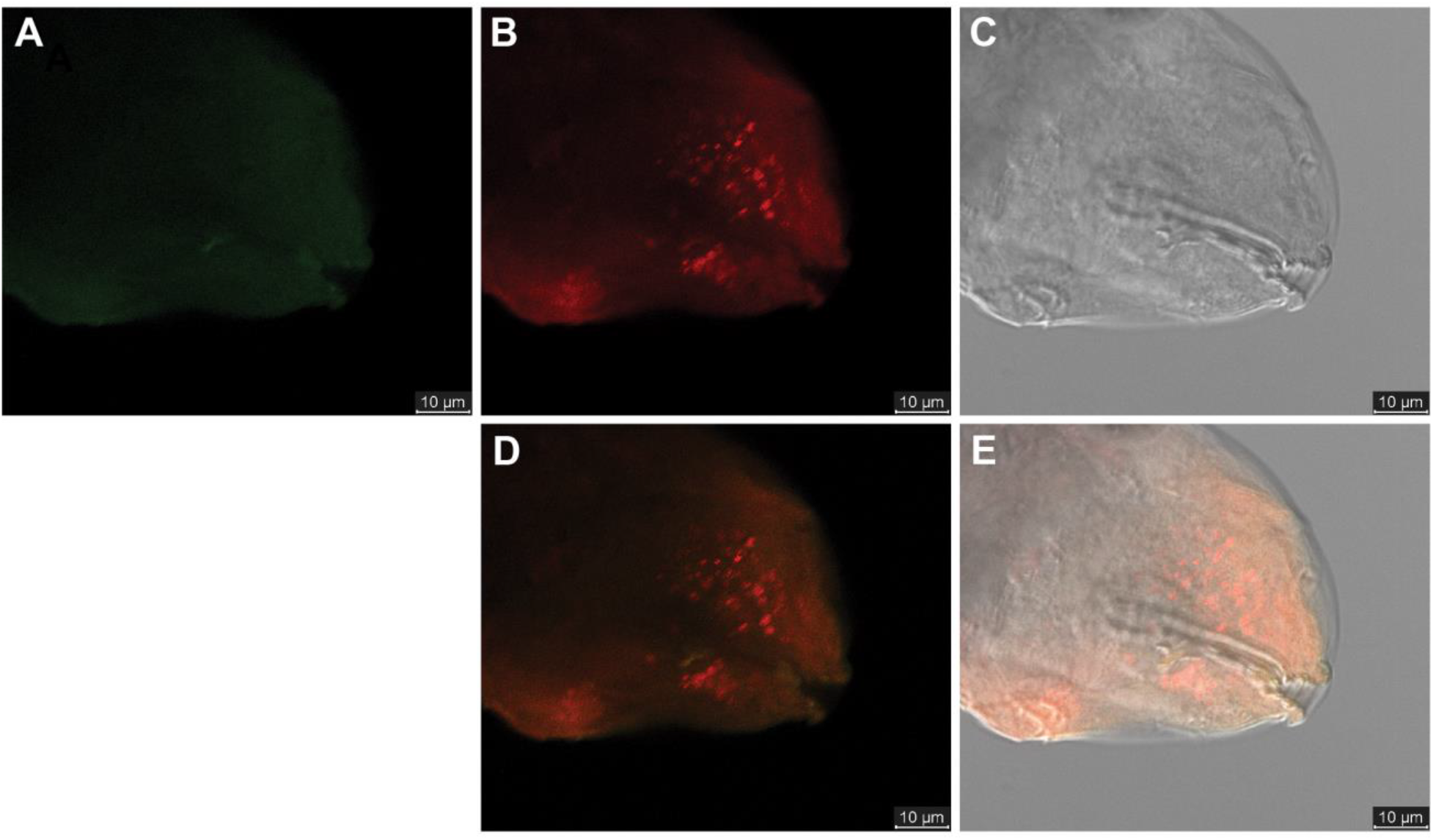
Localization of RickB1 without colocalization of NONEUB strongly indicates the presence of *Rickettsia* in a specimen of *Paramacrobiotus* sp. The specimen was labeled with RickB1 and NONEUB and visualized using confocal microscopy. The viewing field is focused on the head of the specimen. Scale bars = 10 μM, pixel size = 0.106 by 0.106 μm, zoom = 2.69. **(A)** Visualization of the specimen reveals little to no NONEUB signal (green) **(B)** Multiple, ∼1 μm diameter RickB1 signals (red) are visible throughout the head of the specimen. As these do not colocalize with NONEUB, they are likely to be genuine signals from the binding of RickB1 to *Rickettsia* ribosomal RNA **(C)** No structures that may cause autofluorescence are visible through DIC in the location of the RickB1 signals **(D)** Overlay of green and red channels **(E)** Overlay of green and red channels with DIC image.

## Discussion

Contamination is a major concern in microbiome studies of low microbial biomass samples such as tardigrades (Davis et al., 2018; Eisenhofer et al., 2019; Karstens et al., 2019; Tibbs-Cortes et al., 2022). Notably, our recent survey of tardigrade microbiomes identified and removed 986 contaminant taxa using *in silico* methods, including five of what would otherwise have the top ten most abundant bacterial taxa in the study, despite the use of recommended sterile laboratory techniques. These results demonstrate the critical need for careful laboratory and *in silico* mitigation of contamination in order to obtain useful data in tardigrade microbiota surveys (Tibbs-Cortes et al., 2022). In light of these concerns, visualization or other methods to independently verify the presence of bacterial taxa of interest in association with tardigrades will be particularly valuable.

Here, we utilized FISH to generate additional evidence for the presence of *Rickettsia* in tardigrades. Initial experiments utilized the *Rickettsia*-specific probe RickB1 and the general bacterial probe EUB338 to stain specimens. Epifluorescence microscopy revealed that RickB1 colocalized along with EUB338 to circular regions within a specimen of *P. tonollii*, potentially indicating the presence of *Rickettsia* within the specimen. The 1-2 μm points are similar in size and shape to cells of *Rickettsia* species (Socolovschi et al., 2013). The specimen also possessed some areas with EUB338 localization but not RickB1 signals (Figure 1, white arrow), potentially indicating the presence of other bacteria not belonging to the genus *Rickettsia*. Additionally, the absence of similar signals in the red and green channels in the no-probe controls indicates that the EUB338 and RickB1 colocalizations are not false positives caused by autofluorescence of structures in the tardigrade. However, there is some similarity between the signals shown in Figure 1 and the false positive signals observed in Figure 2; therefore, without the use of a nonsense probe, it is difficult to conclude if the RickB1 and EUB338 colocalizations are false positives.

The nonsense probe NONEUB was used to identify regions of autofluorescence and areas of probe accumulation. Tardigrades were hybridized with both NONEUB and RickB1, and specimens were visualized with confocal microscopy. Certain specimens displayed colocalization of both NONEUB and RickB1, demonstrating that the use of a nonsense probe along with the specific probe was capable of detecting false positives (Figure 2). Such false positives could be due to nonspecific incorporation of the two probes or due to autofluorescence of tardigrade tissues. Indeed, some of the regions with both NONEUB and RickB1 corresponded to areas with visible structure under DIC, indicating that these regions might represent autofluorescent anatomical structures in the tardigrade. From this we conclude that nonsense probes are important to eliminating false positives when using FISH in tardigrades.

One sample identified as *Paramacrobiotus* sp. displayed multiple RickB1 signals that did not colocalize with NONEUB (Figure 3). These RickB1 signals were circular with diameter ≤1 μm. Based on the presence of RickB1 signals and the absence of corresponding NONEUB localization, it is likely that these points represent *Rickettsia* within the tardigrade. The shape, size, and pattern of distribution of the putative *Rickettsia* in our specimen are indeed similar to those of *Rickettsia* observed in FISH experiments of other ecdysozoans (Perotti et al., 2006; Pilgrim et al., 2020; Thongprem et al., 2021; Thongprem et al., 2020). This further supports the conclusion that we successfully detected *Rickettsia* in tardigrade tissues. Although the use of a *Rickettsia*-specific probe alone can detect *Rickettsia* in ecdysozoans (Caspi-Fluger et al., 2012; Gottlieb et al., 2006; Kliot et al., 2014; Thongprem et al., 2021; Thongprem et al., 2020), future experiments using a three probe labeling scheme of RickB1 and NONEUB with the addition of EUB338 as a positive control would be useful to confirm our conclusion that *Rickettsia* is present in tardigrades. Because we analyzed tardigrades from the samples with the highest abundance of OTU 180 (classified as *Rickettsia*) from our previous study (Tibbs-Cortes et al., 2022), it is possible that the putative *Rickettsia* observed in our specimen is the same as OUT 180. This could be confirmed by targeted sequencing of the 16S rRNA gene and comparison to the representative sequence of OTU 180 as done by Guidetti et al. (2019b).

The relationship between *Rickettsia* or other endosymbionts and tardigrades has yet to be characterized. Because of their intracellular nature and reliance on vertical transmission, some members of *Rickettsia* and other endosymbiotic taxa have adapted to manipulate host reproduction (McGinn & Lamason, 2021; Moran et al., 2008). It has therefore been hypothesized that tardigrades infected with *Rickettsia* or other endosymbionts may exhibit altered reproductive physiology (Guidetti et al., 2019b; Kaczmarek et al., 2020; Tibbs-Cortes et al., 2022; Vecchi et al., 2018). It is possible that the putative *Rickettsia* identified in this work may affect the reproduction of the host *Paramacrobiotus* sp. While some members of the genus *Paramacrobiotus* exhibit parthenogenesis (Guidetti et al., 2019a; Kaczmarek et al., 2020), others including *P. tonollii* have only been documented as exhibiting bisexual reproduction (Lemloh et al., 2011). Further analysis of the *Paramacrobiotus* found in our collection site would be necessary to determine their reproductive methods and whether infection with *Rickettsia* affected fecundity or reproduction. In addition to vertical transmission, it is also possible that the putative *Rickettsia* identified here may be acquired by tardigrades horizontally between individuals or from environmental sources such as plants (Caspi-Fluger et al., 2012; Chrostek et al., 2017; Moran et al., 2008). Regardless of transmission route, *Rickettsia* infections are known to have a wide variety of effects on hosts, and therefore additional work be necessary to identify and characterize any potential parasitic or mutualistic effects caused by *Rickettsia* on tardigrades (McGinn & Lamason, 2021; Vecchi et al., 2018). Finally, the presence of endosymbiotic taxa in tardigrades raises questions as to how endosymbionts fare during cryptobiosis. If obligate endosymbionts reliant on vertical transmission are present in tardigrades, these would have to survive along with their tardigrade hosts during cryptobiosis in order to be transmitted to offspring. Tardigrade endosymbionts might therefore rely on the cryptobiosis-related molecules of their host to survive cryptobiosis, as suggested by Vecchi et al. (2018).

In addition to providing further evidence for the presence of endosymbiotic bacteria in tardigrades, the methodology presented here may be of additional use in understanding the tardigrade microbiota. For example, it is possible that some putative tardigrade endosymbionts identified by next generation sequencing are actually endosymbionts of the prey of tardigrades (Vecchi et al., 2018). In such cases, localization of FISH probes to the gut contents of tardigrades but not tardigrade tissues would be expected. Additionally, performing FISH on tardigrade embryos could determine if endosymbiotic taxa are maternally inherited in certain tardigrades. Finally, FISH is not limited to the identification of endosymbiotic bacteria. For example, we previously identified putative phytopathogens in the tardigrade community microbiota. However, the amplicon sequencing approach we used lacks sufficient resolution in many cases to distinguish a phytopathogenic isolate from a closely related, innocuous isolate (Tibbs-Cortes et al., 2022). FISH probes designed to target genes encoding phytopathogenic virulence factors could potentially be used to determine if tardigrades carry plant pathogens.

## Conclusion

We utilized FISH to determine if endosymbiotic *Rickettsia* were present within tardigrade tissues. Use of a *Rickettsia*-specific probe, RickB1, and a nonsense probe, NONEUB, allowed us to distinguish between false postives and genuine RickB1 signals. We observed multiple RickB1 signals without NONEUB localization in the head of a specimen of *Paramacrobiotus*. These signals were similar to the signals of *Rickettsia* observed in other ecdysozoa. Therefore, we have provided additional data supporting the presence of endosymbiotic bacteria within tardigrades.

## Acknowledgements

The authors would like to thank Sandra McInnes, Diane Nelson, and Emma Perry for their assistance in tardigrade identification, the Iowa State University Microscopy Facility for training and input with confocal microscopy, the Iowa State University Plant Pathology and Microbiology program for use of the epifluorescence microscope, and the organizers and other attendees of the 15^th^ International Symposium on Tardigrada for a wonderful conference.

